# Continuous constant pH Molecular Dynamics Simulations of Transmembrane Proteins

**DOI:** 10.1101/2020.08.06.239772

**Authors:** Yandong Huang, Jack A. Henderson, Jana Shen

## Abstract

Many membrane channels, transporters, and receptors utilize a pH gradient or proton coupling to drive functionally relevant conformational transitions. Conventional molecular dynamics simulations employ fixed protonation states, thus neglecting the coupling between protonation and conformational equilibria. Here we describe the membrane-enabled hybrid-solvent continuous constant pH molecular dynamics method for capturing atomic details of proton-coupled conformational dynamics of transmembrane proteins. Example protocols from our recent application studies of proton channels and ion/substrate transporters are discussed.

## 1 Introduction

### 1.1 pH-dependent conformational dynamics of membrane proteins

Solution pH plays a critical role in the functions of many transmembrane proteins, including secondary active transporters, proton channels, and G-protein-coupled receptors (GPCRs). For example, pH gradient across the membrane drives the transport of sodium ions through the sodium-proton antiporter NhaA, which is essential for cellular sodium and proton homeostasis in Escherichia coli [54, 60]. Proton channels, such as the voltage-gated proton channel Hv1 [65], conduct protons across the membrane in response to a change in cellular pH [17, 16]. Solution pH also modulates the functions of membrane receptors. For example, the ovarian cancer GPCR1 and the closely related GPR4 regulate cell-mediated responses to acidosis in bone [51].

pH-dependent transmembrane proteins often couple proton sensing with conformational changes [19]. For example, the inward- and outward-facing crystal structures of the sodium-proton antiporter NapA from Thermus thermophilus [46, 14] reveal an elevator-like alternating-access model, although the detailed mechanism of pH-dependent conformational transition remains elusive. In many other cases, however, the conformational change is unknown due to the lack of active-state structure. For example, only inactive crystal structures have been resolved for the aforementioned antiporter NhaA [37, 47] and proton channel Hv1 [68]. Apart from high-resolution structure models, p*K*_*a*_’s of active-site residues are important for mechanistic elucidation, as they inform protonation states at a certain pH, thereby unveiling residues involved in proton binding/release coupled to the conformational transition. However, p*K*_*a*_’s of membrane proteins are extremely challenging if possible at all to experimentally measure. Take NhaA as an example, electrophysiology and biochemical experiments established that it transports two protons in exchange for one sodium above pH 6.5 [67, 61]; however, in the absence of an outward-facing crystal structure and experimental p*K*_*a*_’s, the identities of proton-binding active-site residues remain controversial.

### 1.2 Continuous constant pH molecular dynamics methods

Constant pH molecular dynamics (pHMD) simulations [9, 72, 12] offer a unique means to describe proton-coupled conformational dynamics and calculate p*K*_*a*_ values [72, 76, 3]. Current pHMD methods can be grouped into two classes. In discrete pHMD (DpHMD) methods [5, 55], protonation states of titratable groups are periodically updated using Monte-Carlo sampling following molecular dynamics steps. Rooted in the *λ*-dynamics method for free energy calculations [45], continuous constant-pH molecular dynamics (CpHMD) methods [48, 41, 73, 18, 74, 13, 34, 4, 35, 26, 27] make use of fictitious *λ* particles to sample protonation states simultaneously with atomic coordinates. A *λ* particle carries a mass of a regular heavy atom (e.g., 10 atomic mass units) and the *λ* coordinate continuously evolves between 0 and 1, where 0 and 1 represent protonated and deprotonated states, respectively. The total Hamiltonian includes the kinetic energy of the fictitious particles and the nonbonded term is modified by linearly scaling electrostatic and van der Waals energies according to *λ* coordinates. Thus, CpHMD directly couples conformational dynamics of the protein with protonation equilibria of all ionizable sites at a specified pH. Following a CpHMD simulation at a single pH or a set of replica-exchange CpHMD simulations at multiple pH conditions, the trajectories of *λ* coordinates are extracted, allowing the calculation of p*K*_*a*_’s for all titratable sites.

The first two CpHMD methods [48, 41], implemented in CHARMM molecular dynamics package [6], make use of the generalized Born (GB) implicit-solvent models for both proton titration and conformational dynamics. Since then, several CpHMD methods have been developed and implemented in CHARMM. The hybrid-solvent CpHMD [73] was developed, which takes advantage of accurate description of conformational states in explicit solvent and rapid convergence of protonation state sampling in GB solvent. Additionally, the pH replica-exchange scheme [73] was developed to accelerate convergence of protonation and conformational state sampling. To overcome the limitations of implicit-solvent models, all-atom CpHMD^MS*λ*D^ with force switching was developed that uses explicit solvent to sample both titration and conformational dynamics [26, 27]. To avoid the potential artifacts due to electrostatic cutoff and to further improve accuracy, particle-mesh Ewald (PME) all-atom CpHMD [34] was developed. CpHMD methods have also been implemented in the popular Amber [8] and GROMACS [64] packages, including GBNeck2-CpHMD [35, 28] in Amber and the all-atom CpHMD [18] in GROMACS [64]. Here we focus on the extensively validated [75, 22, 11, 33, 79, 23, 29, 30, 69] pH replica-exchange hybrid-solvent CpHMD method in CHARMM [73], which has been further developed to enable mechanistic studies of membrane proteins [33].

### 1.3 Membrane-enabled hybrid-solvent CpHMD with pH replica-exchange

In the membrane-enabled hybrid-solvent CpHMD method [33], conformational dynamics of a transmembrane protein is sampled in explicit lipids and water, while protonation states are sampled using the membrane GBSW model [38]. Combined with the pH replica-exchange protocol [73], this method allows detailed description of proton-coupled conformational dynamics of transmembrane proteins (Fig. 1). For example, in the simulation study of NhaA [33], a mechanism emerged that explains the pH-dependent activation of NhaA, reconciling a long standing controversy. According to the calculated p*K*_*a*_ values and conformational dynamics [33], Asp164 and Lys300 were identified as proton binding residues, consistent with the recent crystal structures of NhaA [47] and the homologous protein NapA [14]. When NhaA is exposed to the cytoplasmic pH, Asp164 releases the first proton and triggers the hydrophobic gate on the cytoplasm side to open, allowing the entrance of a sodium ion [33]. The sodium ion then binds to three active-site residues, which induces the release of a second proton from Lys300 [33]. The same protocol used to study NhaA has also been applied to investigate the proton-coupled conformational dynamics of the proton channel M2 from influenza virus [11] and the E. coli multi-drug efflux pump AcrB [79]. Most recently, a uniform electric field feature has been added to the membrane-enabled hybrid-solvent CpHMD simulations, allowing us to investigate the activation mechanism of a voltage-gated proton channel Hv1 (Fig. 2).

**Fig. 1.**
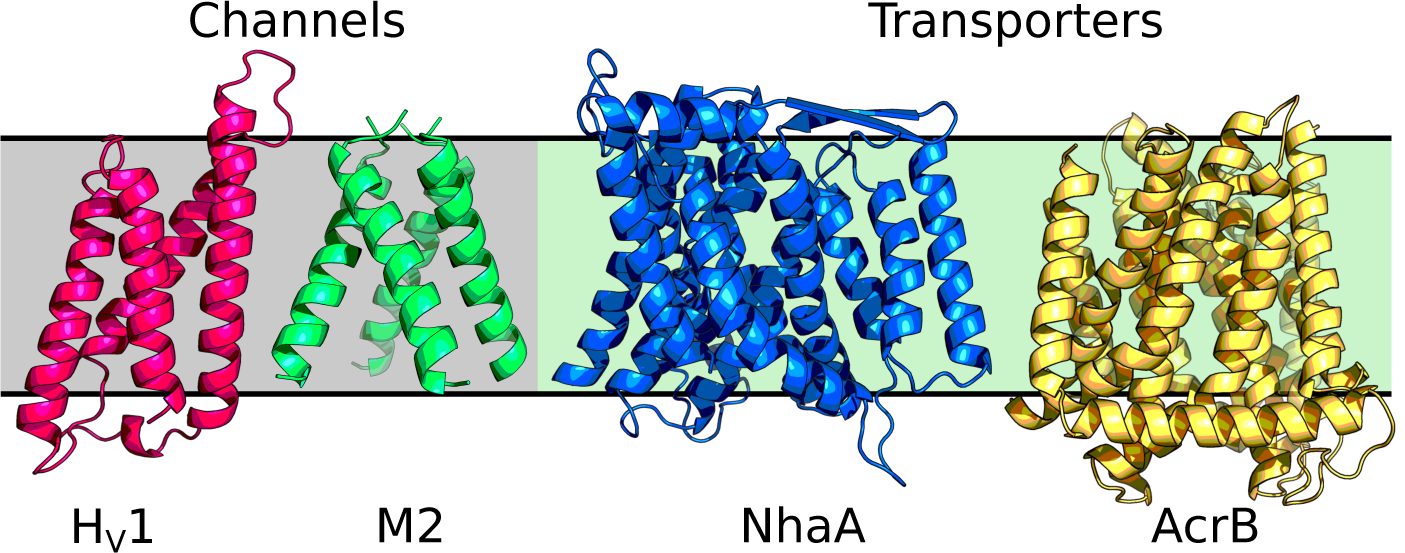
Transmembrane proteins that have been studied by membrane-enabled hybrid-solvent CpHMD simulations with pH replica-exchange [33]: M2 proton channel [11] (PDB: 3LBW [2]), voltage-gated Hv1 proton channel (Henderson and Shen, unpublished data, PDB: 3WKV [68]), sodium-proton antiporter NhaA[33] (PDB: 4AU5 [47]), and multi-drug efflux pump AcrB[79] (PDB: 4DX5 [21]).

**Fig. 2.**
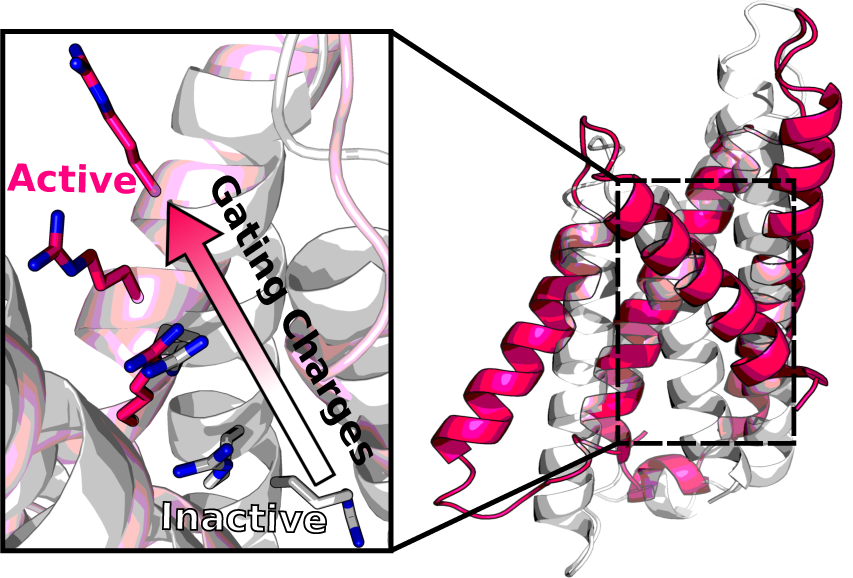
Membrane-enabled hybrid-solvent CpHMD simulations with pH replica-exchange revealed conformational activation of the voltage-gated proton channel Hv1, as indicated by the upward movement of gating charges (Henderson and Shen, unpublished data).

Despite the recent successes of membrane-enabled hybrid-solvent CpHMD simulations (Fig. 1), several important limitations remain. For example, the implicit representation of solvent and bilayer employed in the titration dynamics limits the accuracy of p*K*_*a*_ estimation, particularly for buried active-site residues and those interacting with lipids or small-molecule ligands. In general, p*K*_*a*_ shifts relative to solution (model) p*K*_*a*_ are underestimated for interior residues and overestimated for lipid-interacting ones [33, 11, 79]. Further, explicit interactions with lipids and ions, which may become important for protonation state changes, cannot be accounted for. While PME all-atom CpHMD [34] has the potential to overcome the limitations, our preliminary study of the M2 channel indicated that the calculated p*K*_*a*_’s from all-atom CpHMD have larger systematic errors, likely a result of underpolarization with additive force fields. Thus, the hybrid-solvent CpHMD method remains a method of choice until a solution is found to remedy the force field issue. A future direction is to incorporate polarizable force fields such as the Drude model [49] in PME all-atom CpHMD. Due to the high computational cost of simulations with polarizable force fields, a GPU implementation is desirable.

## 2 Methods

### 2.1 Setting up a CpHMD simulation for transmembrane proteins

Here we discuss the protocol of membrane-enabled hybrid-solvent CpHMD simulations in CHARMM [6], which can be divided into three stages (Fig. 3). In Stage 1, the protein/bilayer complex is constructed and undergoes an initial equilibration. In Stage 2, with the protein structure restrained the lipids are equilibrated to the appropriate structural and phase behavior. In Stage 3, protein sidechains are made titratable and CpHMD is invoked to equilibrate the system at a specified pH condition followed by production replica-exchange CpHMD at multiple pH conditions.

**Fig. 3.**
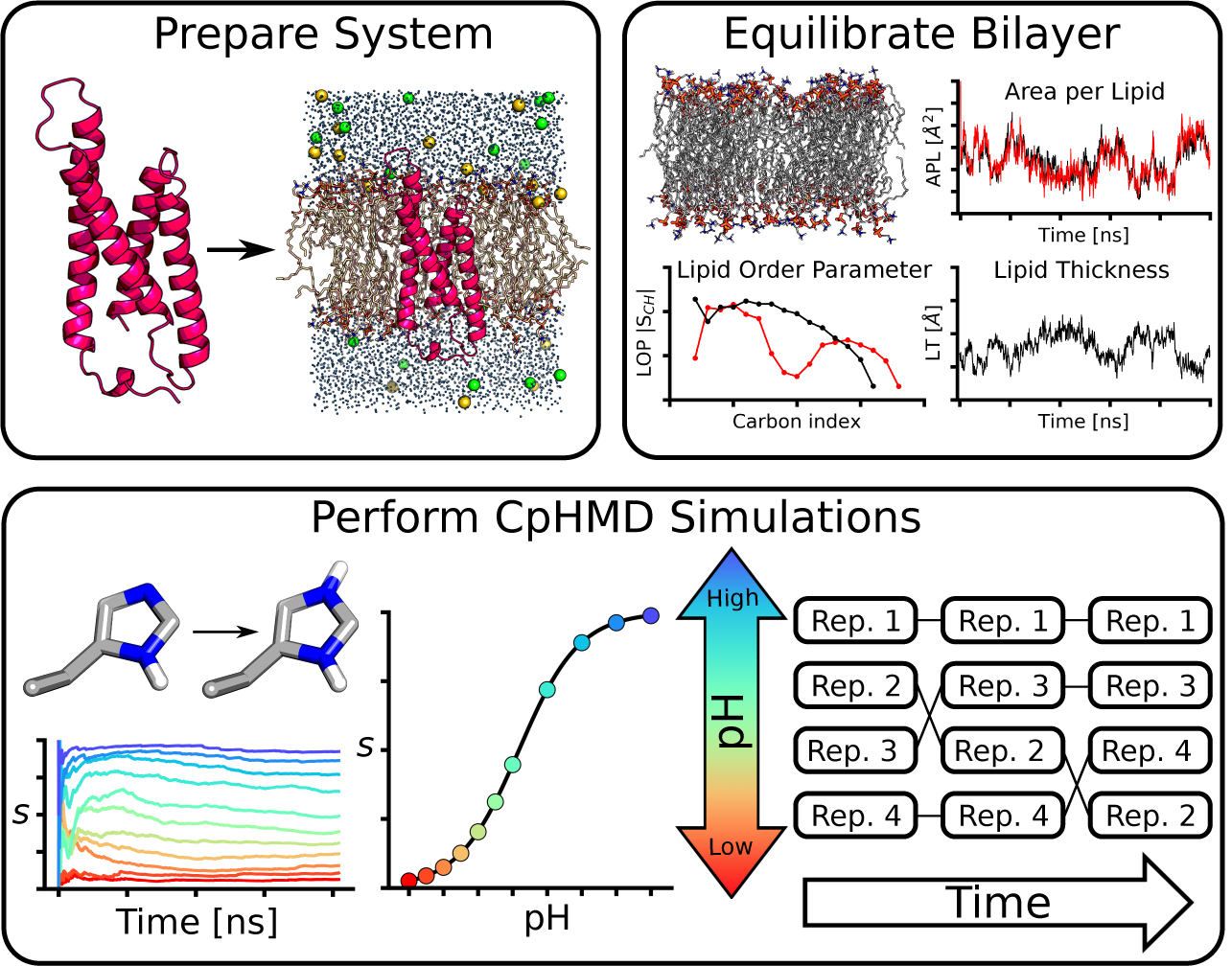
Overview of the three-stage protocol for membrane-enabled hybrid-solvent CpHMD simulations with pH replica-exchange.

**Stage 1. Prepare protein/bilayer complex**. First, we retrieve the protein coordinates from the database, Orientations of Proteins in Membranes [50], which provides coordinates of membrane proteins by orienting the structure with respect to the membrane normal using theoretical calculations and comparisons to experimental data [50]. We then use MMTSB toolset [24] or other tools such as SWISS-MODEL [77] to add or replace missing residues, heavy atoms, or mutation sites. The HBUILD facility [7] in CHARMM is used to add missing hydrogens. Note, in this stage, all titratable sites are assumed to be in the “anticipated” protonation states based on the experimental pH condition and hydrogen bond/salt bridge patterns (for His). If no experimental pH can be found, default protonation states corresponding to physiological pH (7.4) should be used: Asp(-)/Glu(-)/Lys(+)/Arg(+)/Cys(0). Unless His is in a salt bridge, HSE or HSD (based on the hydrogen bonding pattern) should be used.

Since amino acid sequences given in a PDB file are often truncated, we suggest capping the terminal groups of each protein chain: for N-terminus use CH_3_CO (patch ACE), and for C-terminus use NH_2_ (patch CT2) or NHCH_3_ (patch CT3). However, if a terminal group interacts with a part of the protein, the free ionized form, 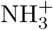 and COO^−^, should be used. Since CpHMD simulation will be conducted at multiple pH in Stage 3, we should consider possible protonation state changes. This consideration is especially relevant for N-amino group, since its solution p*K*_*a*_ is 8.5 [59], only one unit above the physiological pH. If N-amino group will be allowed to titrate, make sure the model parameters are included in the PHMD parameter file.

Once the protein structure is complete, a very short minimization in implicit solvent (e.g., with the GBSW model [39]) may be conducted with the steepest descent (SD) and adopted basis Newton-Raphson (ABNR) algorithms to relax unfavorable positions of hydrogens. It is important that during the minimization, protein heavy atoms should be fixed or restrained. For example, two rounds of 50 steps of SD followed by 10 steps of ABNR are performed, first with heavy atoms fixed and then with a harmonic force constant of 5 kcal/mol/Å^2^. If an extensive number of atoms have been added we also suggest that these sites be energy minimized, but the overall structure should be kept as close as possible to the crystal structure. If crystal waters are resolved in the structure these should be retained.

The rest of Stage 1 follows the CHARMM-GUI protocol [40] for preparing a protein/membrane complex for MD simulations. The protein is placed in a lipid bilayer with water above and below the bilayer. Counterions for neutralizing the system and additional ions to reach the specified ionic strength are added. The assembled protein/membrane complex undergoes a series of initial equilibration steps (Table 1, Stage 1) to relax the lipid positions before subjecting to a lengthy membrane equilibration step in Stage 2.

**Table 1.** Example equilibration steps in all three stages

| Step | Ensemble | Timestep<br>(fs) | Length<br>(ns) | Restraint force constant<br>(kcal/mol/Å<sup>2</sup>) |  |  |  |
| --- | --- | --- | --- | --- | --- | --- | --- |
| **Stage 1. Initial equilibration** |  |  |  | BB<sup>a</sup> | SC+XWat<sup>a</sup> | Lipid<sup>b</sup> | BWat<sup>c</sup> |
| 1 | NVT | 1 | 0.05 | 10.0 | 5.0 | 2.5 | 2.5 |
| 2 | NVT | 2 | 0.05 | 5.0 | 5.0 | 2.5 | 2.5 |
| 3 | NPT | 2 | 0.1 | 5.0 | 5.0 | 1.0 | 1.0 |
| 4 | NPT | 2 | 0.1 | 5.0 | 2.5 | 0.5 | 0.5 |
| 5 | NPT | 2 | 0.1 | 2.5 | 2.5 | 0.1 | 0.1 |
| 6 | NPT | 2 | 0.1 | 2.5 | 2.5 | 0.0 | 0.0 |
| **Stage 2. Membrane equilibration** |  |  |  | BB + SC |  |  |  |
| 1 | NVT | 2 | 0.2 | 2.5 |  |  |  |
| 2 | NPT | 2 | 100 | 2.5 |  |  |  |
| **Stage 3. CpHMD equilibration** |  |  |  | BB + SC |  |  |  |
| 1 | NVT | 2 | 0.1 | 1.0 |  |  |  |
| 2 | NPT | 2 | 0.1 | 1.0 |  |  |  |
| 3 | NPT | 2 | 0.8 | 0.0 |  |  |  |

<sup>a</sup> BB and SC refer to protein backbone and side chain heavy atoms, respectively. XWat and BWat refer to crystallographic and bulk water oxygen atoms, respectively. <sup>b</sup> Lipid heavy atoms are restrained with a planar harmonic potential to ensure that lipid tails are close to the bilayer center ( $-5 \text{ Å} < Z < 5 \text{ Å}$ ), and lipid head groups are close to the bilayer surface ( $Z = \pm 19 \text{ Å}$ for POPE). <sup>c</sup> A planar harmonic restraint is applied to the bulk water oxygen atoms to keep them away from the membrane hydrophobic region.

**Stage 2. Equilibrate bilayer**. Once a protein/membrane complex is assembled and relaxed, the lipid bilayer needs to be fully equilibrated before protein conformational states can be sampled. Simulation in this stage can be carried out using any scalable simulation engine, such as GROMACS [1], NAMD [63], or OpenMM [20]. Typically, a 0.2-ns restrained MD is performed in the NVT ensemble followed by a 100-ns restrained MD in the NPT ensemble (see Table 1, Stage 2). During the equilibration, protein backbone and side chain heavy atoms are restrained with a harmonic force constant of 1.0 kcal/mol/Å^2^.

Surface area per lipid, bilayer thickness, and lipid order parameters are quantities commonly used to assess the phase behavior and fine structure of the simulated bilayer [43]. A lipid bilayer is considered equilibrated once all three quantities are converged to experimental values for the specified lipid type or those obtained in the force field validation study [43]. The program VTMC [56] can be used to calculate surface area per lipid; it excludes lipids within 5 Å from the protein, as protein-lipid interactions can artificially decrease the value. Bilayer thickness here refers to the hydrophobic thickness, which is the distance between the average *Z*-positions of the lipid C2 atoms from the top and bottom leaflets. Lipid order parameters *S*_CH_ indicate the rigidity of the lipid tails [71] and are defined as

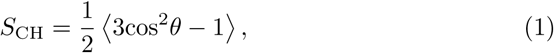

where *θ* is the angle between the C-H bond of the lipid tail methylene and the membrane normal (*z*), and the angular brackets denote ensemble average. *S*_CH_ can be calculated using a Tcl script for VMD [36].

**Stage 3. Perform CpHMD titration simulations**. Following equilibration of bilayer, the system can be further equilibrated using membrane-enabled [33] hybrid-solvent CpHMD simulation at a single pH. From the final snap shot of Stage 2, dummy protons need to be added to all Asp, Glu, and His sidechains. For Asp and Glu, dummy protons are placed in the *syn* position. Note, the dihedral energy barrier to the rotation around the carboxylic bond is raised (to 3 kcal/mol) in the force field [41] to prevent a dummy hydrogen from moving to the less favorable *anti* position where it becomes unlikely to protonate (“ghost” proton) [55, 41]. For His, a proton is added to either *δ* or *E* nitrogen to form a doubly protonated HSP. Addition of dummy hydrogens requires a brief retrained energy minimization, e.g., two rounds of 50 steps of SD followed by 10 steps of ABNR. In the first round all heavy atoms are fixed in positions, and in the second round a harmonic restraint of 5.0 kcal/mol/Å^2^ is applied to them. Next, three short restrained CpHMD simulations are performed at crystallization or physiological pH (see Table 1, Stage 3). The first and second CpHMD equilibration steps are run for 0.1 ns in the NVT and NPT ensembles, respectively, with protein heavy atoms restrained with a harmonic force constant of 1.0 kcal/mol/Å^2^. The third unrestrained equilibration step is run in the NPT ensemble for 0.8 ns. To prevent lateral drift of the protein, a cylindrical restraint with a harmonic force constant of 0.1 kcal/mol/Å^2^ is applied using the MMFP facility in CHARMM [6].

Following equilibration at a single pH, production CpHMD simulations can begin using the pH replica-exchange protocol [73]. First, a simulation pH range is selected by considering the active pH range of the protein or the one that represents experimental or physiological pH conditions. The pH replicas need to be placed close enough to obtain sufficient exchange probabilities (e.g., *>* 20%), which requires overlap between the potential energy distributions of neighboring replicas. Typically, a 0.5 pH unit increment provides desired exchange probabilities. Each replica is simulated in the NPT ensemble, with exchange attempts occurring every 500 or 1000 MD steps between adjacent replicas. We typically observe an average exchange rate of at least 40%. More replicas at smaller intervals (e.g., 0.25 pH units) may be added to increase the exchange rate. The effectiveness of sampling enhancement can be tracked by monitoring the movement of replicas through pH space as simulation proceeds. At least several replicas should move across the entire pH ladder to achieve enhanced sampling.

#### Settings in molecular dynamics

Protein is represented by CHARMM22/CMAP force field [52, 53], lipids and ions are represented by CHARMM36 force field [43], and water is represented by CHARMM modified TIP3P model. van der Waals energies are calculated using a switching function from 10 to 12 Å with a cutoff at 14 Å, consistent with the protocol used in the development of CHARMM36 lipid force field [43]. Simulations are conducted under periodic boundary conditions with the PME method [15] for long-range electrostatic energies and forces. The real space calculation uses a 12-Å cutoff and the reciprocal space calculation uses a 1-Å grid spacing and 6th-order spline interpolation. The nonbond neighbor list is heuristically updated in CHARMM [6] or every 10 MD steps in GROMACS [64].

To allow for a 2-fs integration time step, all bonds involving hydrogens are constrained using SHAKE [66] in CHARMM [6], LINCS [31] in GROMACS [64], or rigidBonds in NAMD [63]. In NPT simulations, temperature is maintained at 310 K by Nóse-Hoover thermostat [58, 32] and pressure is maintained at 1 atm by Langevin piston pressure-coupling algorithm in CHARMM [25] or Parrinello-Rahman barostat [62] in GROMACS.

#### CpHMD specific settings

Unless otherwise noted, default CpHMD settings should be used. In membrane-enable hybrid-solvent CpHMD simulations [33], the protein is centered in the lipid bilayer and an exclusion cylinder, e.g., with a radius of 25 Å, is placed along the membrane normal to exclude the low dielectric slab used in the implicit membrane model [38] from influencing the titration of interior residues in the protein. The cylinder should encompass the protein with no or minimal overlap with lipids. A center of mass restraint is applied to the protein with a harmonic force constant of 1.0 kcal/mol/Å^2^ via the MMFP facility in CHARMM[6]. The thickness of the low-dielectric slab is set to the hydrophobic thickness of the lipid bilayer, defined as the difference between the average *z* positions of lipid C2 atoms from the top and bottom leaflets, e.g., about 35 Å for POPE. The half of membrane switching length [38] for the dielectric transition region between the low-dielectric slab and bulk water is set to 2.5 Å. The default GBSW atomic input radii [57, 10] are used for protein. For small molecules, the generic atomic radii in the GBSW radii file or van der Waals radii in the force field [44] should be used. Titration (*λ*) and tautomeric state (*x*) coordinates are updated every 5 MD steps (default for hybrid-solvent CpHMD), allowing for water and lipid relaxation [73]. Langevin dynamics with a friction coefficient of 5 ps^−1^ is used (default for CpHMD). The mass of *λ* and *x* particles are set to 10 atomic mass units (default for CpHMD).

### 2.2 p*K_a_* calculations

To calculate the p*K*_*a*_ of a titratable site, the deprotonated fraction at each pH, *S*(pH), is obtained and fit to the generalized Henderson-Hasselbalch equation (Hill equation).

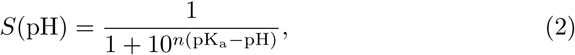

where *n* is the Hill coefficient representing the slope of the transition region in the titration curve. Deviation of *n* from 1 indicates the extent of cooperative (*n >* 1) or anticooperative (*n <* 1) interactions between two or more titrating sites [72]. The deprotonated fraction of a titratable site is calculated as the fraction of time *λ* samples the deprotonated state,

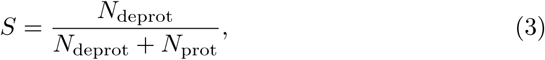

where deprotonated and protonated states are defined as *λ >* 0.8 and *λ <* 0.2, respectively.

For coupled titrating sites, i.e. titration of one site is affected by the protonation state of the other, apparent or macroscopic sequential p*K*_*a*_s are of interest [28]. In the statistical mechanics formulation, the two sequential p*K*_*a*_’s, p*K*_1_ and p*K*_2_, are obtained by fitting the total number of protons bound to the two sites, *N*_prot_, to the coupled titration model below [70, 74].

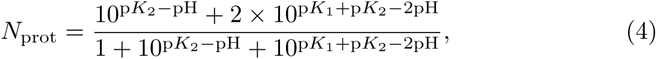

Alternatively, the total deprotonated fraction of the two sites can be fit to the following uncoupled model [78, 27],

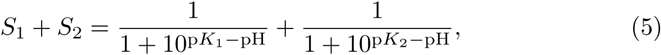

where *S*_1_ and *S*_2_ are the deprotonated fractions of the two sites. p*K*_1_ and p*K*_2_ are the two uncoupled p*K*_*a*_’s.

## Acknowledgement

The authors acknowledge National Institutes of Health (R01GM098818 and R01GM118772) for funding.

## 3 Checklist of Items to be Sent to Volume Editors

Here is a checklist of everything the volume editor requires from you:

□ The final LATEX source files
□ A final PDF file
□ A copyright form, signed by one author on behalf of all of the authors of the paper.
□ A readme giving the name and email address of the corresponding author.

